# Piecemeal regulation of convergent neuronal lineages by bHLH transcription factors in *C. elegans*

**DOI:** 10.1101/2021.03.31.437939

**Authors:** Neda Masoudi, Eviatar Yemini, Ralf Schnabel, Oliver Hobert

## Abstract

Classic cell lineage studies in the nematode *Caenorhabditis elegans* as well as recent lineage tracing in vertebrates have shown that cells of the same type can be generated by distinct cellular lineages that originate in different parts of the developing embryo (“lineage convergence”). Several *C. elegans* neuron classes composed of left/right or radially symmetric class members display such lineage convergence, in that individual neurons of the same class derive from distinct, non-bilaterally symmetric lineages. We show here that the *C. elegans* Atonal homolog *lin-32/Ato*, a bHLH transcription factor, is differentially expressed in neuronal lineages that give rise to left/right or radially symmetric class members. Loss of *lin-32/Ato* results in the selective loss of the expression of panneuronal markers and terminal selector-type transcription factors that confer neuron class-specific features. We discovered that another bHLH transcription factor, the Achaete Scute-homolog *hlh-14* is expressed in mirror image pattern to *lin-32/Ato* in a subset of the left/right symmetric neuron pairs and is required to induce neuronal identity and terminal selector expression on the contralateral side of the animal. These findings demonstrate that distinct lineage histories converge via distinct bHLH factors on the level of induction of terminal selector identity determinants, which thus serve as integrators of distinct lineage histories. We also describe neuron-to-neuron identity transformations in *lin-32/Ato* mutants, which we propose to also be the result of misregulation of terminal selector gene expression.

## INTRODUCTION

The completely elucidated lineage history of every individual cell in the nematode *C. elegans* provides an excellent opportunity to study how lineage affects cellular identity (Sulston and Horvitz 1977; Sulston *et al*. 1983). One intriguing revelation of the lineage description is that phenotypically similar cells can have different lineage histories. This is particularly evident in the nervous system, composed of 302 neurons in the hermaphrodite. Based on anatomy, function and molecular profiles, these 302 neurons can be grouped into 118 different classes, with members of each class being phenotypically similar and often completely indistinguishable by any known criterion (White *et al*. 1986; Hobert *et al*. 2016; Taylor *et al*. 2020). Members of individual neuron classes can have very similar lineage histories. For example, many ventral cord motor neuron classes are composed of members with similar lineage histories (Sulston *et al*. 1983). Also, many classes of bilaterally symmetric neuron pairs are composed of two members whose developmental history is very similar (Sulston *et al*. 1983). However, member of the same neuron class can also very distinct lineage histories. For example, the four bilaterally symmetric cephalic CEP sensory neurons, composed of a bilaterally symmetric ventral neuron pair (CEPV left and CEPV right) and a bilaterally symmetric dorsal neuron pair (CEPD left and right) share similar overall morphology, similar patterns of synaptic connectivity (White *et al*. 1986) and similar molecular composition (Taylor *et al*. 2020). However, the CEPD pair and CEPV pair come from different neuroblast lineages (Sulston *et al*. 1983). Members of other radially-symmetric neuron classes are also phenotypically indistinguishable, despite distinct lineage histories; for example, the IL1, IL2, and OLQ neuron classes are composed of ventral and dorsal pairs which each have distinct lineage histories (Sulston *et al*. 1983). Even neuron classes that are only composed of two bilaterally symmetric can be defined by two lineally distinct neurons. For example, the left and right ASE neuron derive from different embryonic blastomere, ABa and ABp (Sulston *et al*. 1983). Thus, distinct lineage histories can converge onto similar neuronal identities. Similar lineage convergence phenomena have recently been observed in vertebrates as well, including vertebrates (McKenna *et al*. 2016; Wagner *et al*. 2018; Cao *et al*. 2019; Chan *et al*. 2019). For example, specific types of excitatory and inhibitory neurons of the mouse CNS develop through multiple, convergent trajectories(Cao *et al*. 2019).

How is such convergence achieved? Based on past studies of neuronal identity control, one point of convergence of distinct lineage histories are terminal selector transcription factors – post-mitotically expressed master regulators of neuron identity (Hobert 2016). For example, the two lineally distinct ASE neurons both eventually turn on the terminal selector CHE-1, which instructs ASE neuron identity. Similarly, all six members of the IL2 neuron class, despite their distinct lineage histories, co-express the terminal selectors *unc-86, sox-2* and *cfi-1*, which cooperate to control the expression of IL2 identity features (Shaham and Bargmann 2002; Zhang *et al*. 2014; Vidal *et al*. 2015). Similarly, the left and right CAN neurons, which display non-symmetric, distinct lineage histories, coexpress the *ceh-10* homeobox gene, required to specify the identity of both of these neurons (Forrester *et al*. 1998; Wenick and Hobert 2004). These observations indicate that different lineage histories converge onto the expression of similar terminal selectors and that terminal selectors are therefore integrators of distinct lineage histories. What then are the molecular factors that converge in a lineagespecific manner to drive individual terminal selectors in a neuron type-specific manner?

Recent scRNA analysis of developing embryos has revealed molecular correlates to distinct lineage histories, such that the precursors of cells with similar terminal identities were shown to display distinct gene expression profiles, exactly as predicted by their distinct lineage histories (Sulston *et al*. 1983; Packer *et al*. 2019). However, no specific genetic factors have been shown so far to differentially control the identity of cells that converge on similar cell-fate outcomes. In this paper, we show that the *C. elegans* Atonal homolog *lin-32/Ato* (Zhao and Emmons 1995) is differentially expressed in distinct lineages of cells that phenotypically converge on the same neuronal cell classes and, henceforth, selectively affects the specification of only some class members. Moreover, we discovered that the Achaete Scute homolog *hlh-14* (Frank et al. 2003) displays the mirror image expression to *lin-32/Ato* and functions on the contralateral side of the animal.

Our analysis of bHLH transcription factors was originally motivated by our quest to understand how the expression of terminal selector transcription factors is controlled. Terminal selectors have emerged as key regulators of neuronal identity throughout the entire nervous system (Hobert 2016), yet we understand little about how their expression is induced in the embryo (Bertrand and Hobert 2009; Murgan *et al*. 2015). Based on a previously reported effect of *lin-32/Ato* on the expression of the terminal selectors *unc-86* (Baumeister et al. 1996) and *mec-3* (Way et al., 1992), we sought to investigate the role of *lin-32/Ato* further. In a number of different organisms, from flies to worms to vertebrates, Atonal orthologs have mostly been characterized for the proneural activity that imposes neuronal identity on neuroectodermal progenitor cells (Bertrand *et al*. 2002; Baker and Brown 2018). Loss of such proneural activity results in a conversion from a neuronal to an ectodermal skin cell fate. Such proneural functions have been defined for *lin-32/Ato* in the context of number of peripheral sense organs in *C. elegans*, including the postdeirid lineage, the Q lineage, and ray lineages in the male tail (Chalfie and Au 1989; Zhao and Emmons 1995; Portman and Emmons 2000; Zhu *et al*. 2014). These proneural functions are evidenced by lineage changes in which neuroblasts that normally divide to generate multiple distinct neuron types instead convert to skin cells, leading to a failure to generate a number of neuron types (Chalfie and Au 1989; Zhao and Emmons 1995; Portman and Emmons 2000; Zhu *et al*. 2014). However, other effects on neural lineages have been observed in *lin-32/Ato* mutants as well. For example, after its proneural role early in lineage development, *lin-32/Ato* has also been implicated in controlling later aspects of neuronal differentiation in the Q and ray lineages (Portman and Emmons 2000; Miller and Portman 2011; Zhu *et al*. 2014). Cell identity fate transformations are also evident in a subset of dopaminergic neuronproducing lineages (Doitsidou *et al*. 2008) as well as in glia cells (Zhang *et al*. 2020). *lin-32/Ato* has also been shown to affect the differentiation and/or function of a number of additional neuron types, including oxygen sensory neurons (Rojo Romanos *et al*. 2017), touch receptor neurons (Mitani *et al*. 1993), anterior ganglion neurons (Baumeister *et al*. 1996; Shaham and Bargmann 2002), and the AIB interneuron (Hori *et al*. 2018). However, it was left unclear whether these defects are proneural lineage defects, cell-identity transformation defects, or some combination of both these effects.

Here, to facilitate a comprehensive analysis of *lin-32/Ato* function, we begin by describing the expression pattern of *gfp-tagged* LIN-32/Ato. While the expression of *lin-32::gfp* is consistent with a number of previously described functions of *lin-32/Ato*, we identified sites of expression that led us to explore novel functions of *lin-32/Ato*. One consistent theme of *lin-32/Ato* function is its effect on the expression of terminal selector-type transcription factors. Our findings for *lin-32*, as well as another bHLH family of the Atonal superfamily, *hlh-14*, provide novel insights into the molecular basis for the convergence of distinct lineage history onto similar cellular differentiation events, via the regulation of terminal selector expression.

## MATERIAL AND METHODS

### Strains

Strains were maintained by standard methods (Brenner 1974). Previously described strains used in this study are as follow.

Mutant alleles *lin-32(tm1446)* (Doitsidou et al. 2008), *hlh-14(tm295)* (Poole *et al*. 2011).

#### Reporter alleles

*lin-11(ot958[lin-11::gfp::FLAG])* (Reilly et al. 2020), *ceh-32(ot1040[ceh-32::gfp])* (kindly provided by Cyril Cros), *unc-86(ot879[unc-86::nNeonGreen])* (Serrano-Saiz et al. 2018), *unc-42(ot986[unc-42::gfp])(BERGHOFF* et al. 2021).

#### Reporter transgenes

*nIs394 (ngn-1::gfp)* (Nakano et al. 2010), *otIs339 (ceh-43^fosmid^::gfp*) (Doitsidou et al. 2013), *otIs703(Is[flp-3::mCherry]); myIs13 (Is[klp-6::gfp])* (kindly provided by Maryam Majeed), NeuroPAL (*otIs669*)(Yemini et al. 2021), *otIs356 (Is[rab-3::NLS::tagRFP])(SERRANO-SAiz* et al. 2013), *lqIs4 [ceh-10p::GFP + lin-15(n765)] X* (kindly provided by Erik Lundquist), UL1692: *unc-119(ed3); IeEx1692 (Ex[hlh-34::gfp, unc-119(+)])* (Cunningham et al. 2012).

Fosmid-based reporters for *hlh-3, hlh-14, lin-32* and *cnd-1* were generated by insertion of *gfp* at the 3’end of their respective loci using fosmid recombineering (Sarov *et al*. 2012). Fosmid names are indicated in the figures. All fosmids were injected as complex arrays at the concentration of 20 ng/μl with 5ng/μl of *ttx-3::mCherry* as a co-injection marker and up to 85ng/μl of OP50 DNA into N2 and then chromosomally integrated. The array names are: *otIs594 (lin-32^fosmid^::gfp), otIs648 (hlh-3^fosmid^::gfp), otIs713 (hlh-14^fosmd^::gfp). otIs813 (cnd-1^fosmid^::gfp)*

### Microscopy

For fluorescence microscopy, worms were paralyzed by 25mM sodium azide (NaN3) and mounted on 5% agarose pad on glass slides. Images were acquired using an axioscope (Zeiss, AXIO Imager Z.2) or LSM 800 laser point scanning confocal microscope. Representative images are max-projection of Z-stacks. Image reconstruction was performed using Fiji software (Schindelin *et al*. 2012).

4D microscopy and SIMI BioCell (Schnabel *et al*. 1997) was used, as previously described, to analyze embryonic lineage defects of mutant animals as well as bHLH fosmid reporter expression pattern during embryogenesis. Briefly, gravid adults were dissected on glass slide and a single two-cell stage embryo was mounted and recorded over 8 hours of embryonic development. Nomarski stacks were taken every 30 s and embryos were illuminated with LED fluorescence light (470 nm) at set time points during development. The recording was done with Zeiss Imager Z1 compound microscope, using the 4D microscopy software Steuerprg (Caenotec).

## RESULTS

### Embryonic expression pattern of *lin-32/Ato*

Even tough *lin-32/Ato* was identified more than a quarter century ago (Zhao and Emmons 1995), its embryonic expression pattern has not been reported so far. We analyzed a transgenic strain carrying a fosmid in which the *lin-32/Ato* locus has been 3’-terminally tagged with *gfp*, as well as as CRISPR/Cas9-engineered reporter allele, in which *gfp* was also inserted at the 3’ end of the gene (kindly provided by the Greenwald lab), both of which yielded similar expression patterns. Embryonic expression was analyzed using 4D microscopy (Schnabel *et al*. 1997). We focused on the two lineages that produce all but two (PVR, DVC) of the 302 neurons of the nervous system, the AB and MS blastomeres. We Fig.9 that *lin-32/Ato* expresses in a number of different neuronal lineages in the AB lineage, but not the MS lineage, which produces only pharyngeal neurons (**Fig.1**). The earliest expression was observed at the 128-cell stage, in the daughters of the ABalapa blast cells. In other lineages the onset of *lin-32/Ato* expression occurred shortly thereafter while, elsewhere, it commenced as late as a mother neuroblast that generates two terminally-differentiating daughters. In several cases we were not able to record the terminal division of a neuroblast due to the movement of embryos in the egg shell and we may therefore have missed expression in postmitotic cells. If postmitotic expression were to exist, it would be transient since we observed no expression of *lin-32/Ato* in any embryonically-generated neuron in first larval or later stage animals.

**Fig.1:**
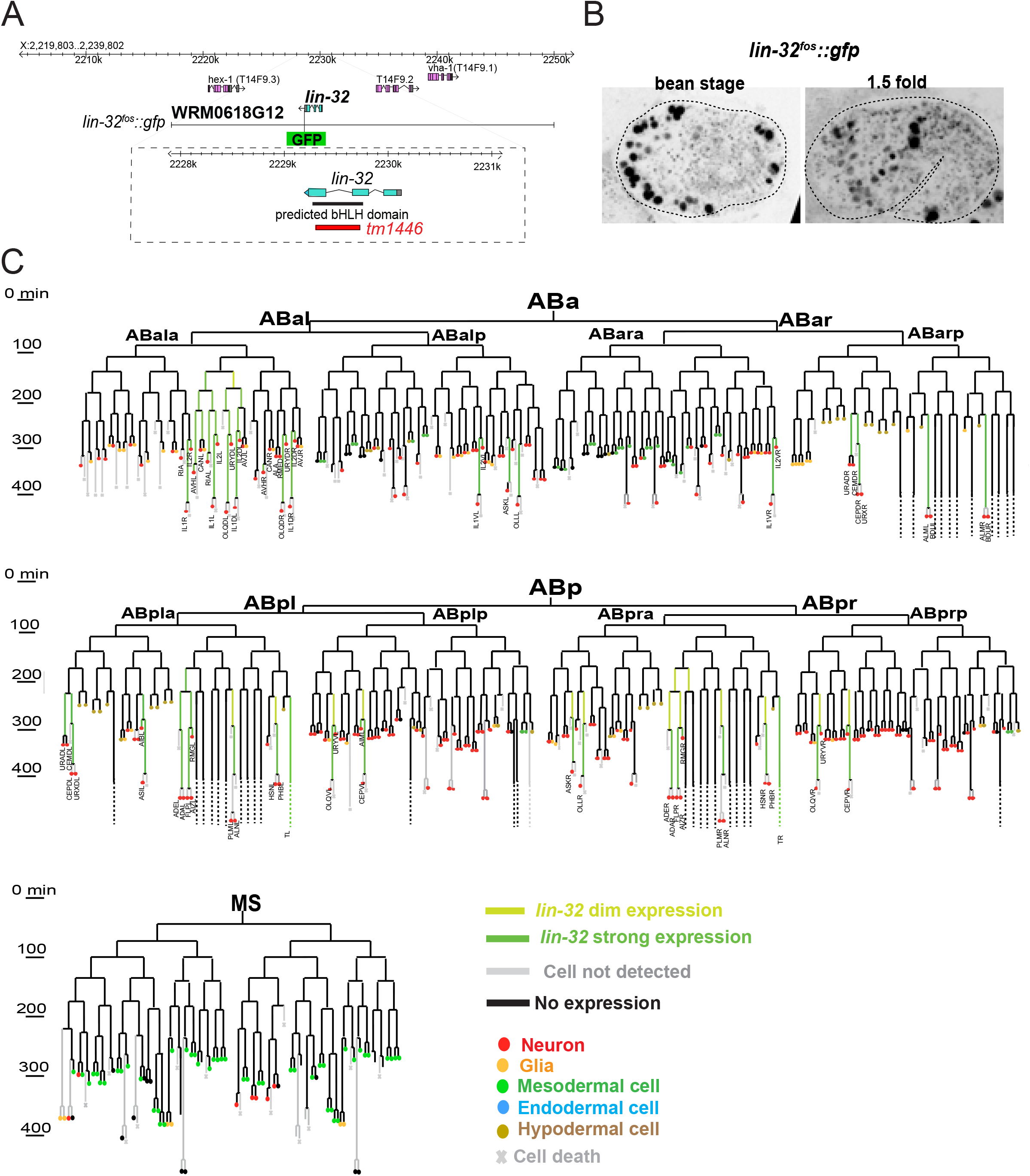
*lin-32/Ato* expression pattern. **A:** Schematic of gene structure shows that the deletion allele *tm1664* removes the bHLH domain from *lin-32/Ato* locus. **B:** Representative images of *lin-32/Ato* fosmid gene expression (*otIs594*) at embryonic stages when most terminal neurons are born. **C:** *lin-32/Ato* fosmid expression (*otIs594*) in the AB and MS lineages, which produce all but 2 of the 118 neuron classes of the hermaphrodite. The *lin-32/Ato* fosmid reporter is first detected shortly after 100 minutes into development in ABalap descendants. Our analysis reveal that *lin-32::gfp* is expressed in both mitotically active neuroblasts during embryogenesis and a subset of postmitotic neurons.

Consistent with expression of Atonal homologs in other organisms (Bertrand *et al*. 2002; Jarman and Groves 2013), we note that the vast majority of lineages that express *lin-32/Ato* produce sensory neurons. Several aspects of the embryonic expression of *lin-32/Ato* are in agreement with previously reported *lin-32/Ato* mutant phenotypes. For example, we observe *lin-32/Ato* expression (**Fig.1**) in lineages that give rise to sensory neurons in the anterior ganglion, consistent with previous studies that reported *lin-32/Ato* to control expression of the *unc-86* terminal selector (Baumeister *et al*. 1996; Shaham and Bargmann 2002). Similarly, we observed expression in lineages that give rise to the URX and CEPD neurons, and in lineages that give rise to the AIB neurons, which are all neurons wherein differentiation defects have been observed in *lin-32/Ato* mutants (Doitsidou *et al*. 2008; Rojo Romanos *et al*. 2017; Hori *et al*. 2018).

### Proneural functions of *lin-32/Ato*

Previous work had shown that *lin-32/Ato* has proneural functions in a number of postembryonically generated sensory neuronal cell types, including the sex-shared postdeirid lineage and the Q-lineage, as well as in male-specific neuronal lineages (Chalfie and Au 1989; Zhao and Emmons 1995; Portman and Emmons 2000). One common feature in these lineages appears to be that the loss of *lin-32/Ato* results in obvious lineage patterning defects, in which normally dividing neuroblasts transform into hypodermal cells. The embryonic expression of *lin-32/Ato* that we described above prompted us to ask whether such neuroblast-to-hypodermal conversions are also observed in embryonically generated neurons of *lin-32/Ato* null mutant animals (*tm1449* deletion allele; **Fig.1**). Nomarksi optics-based lineage tracing of embryonic cell lineages was done using 4D microscopy with SIMI Biocell (Schnabel *et al*. 1997), until 300 minutes of embryonic development. This analysis revealed no obvious cell-division defects or transformations into hypodermal fates (as would be evidenced by changes in nuclear morphology and migratory patterns) in any of the lineages that normally express *lin-32/Ato*.

For a more granular assessment of cell fate, we examined the expression of different sets of panneuronal cell-fate markers. First, we used a nuclear, localized reporter transgene for the panneuronal *rab-3* gene (Stefanakis *et al*. 2015). We counted the number of nuclei and, with the exception of the ventral nerve cord, observed a reduction in overall panneuronal gene expression throughout embryonically generated head and tail ganglia (**Fig.2A**). To further confirm this observation, we used a hybrid reporter construct in which the *cis*-regulatory elements of several panneuronally expressed genes are fused with one another, the “UPN” driver (Yemini *et al*. 2021). We observed the expression of this panneuronal marker gene expression to also be lost in many different normally *lin-32/Ato* expressing lineages (**Fig.2B,Fig.3**). Lastly, we also examined the existence of subnuclear granules (“NUN bodies”), another panneuronal identity features (Pham *et al*. 2021). We observed a reduction of the number of cells with this subnuclear structure (**Fig.2C**).

**Fig.2:**
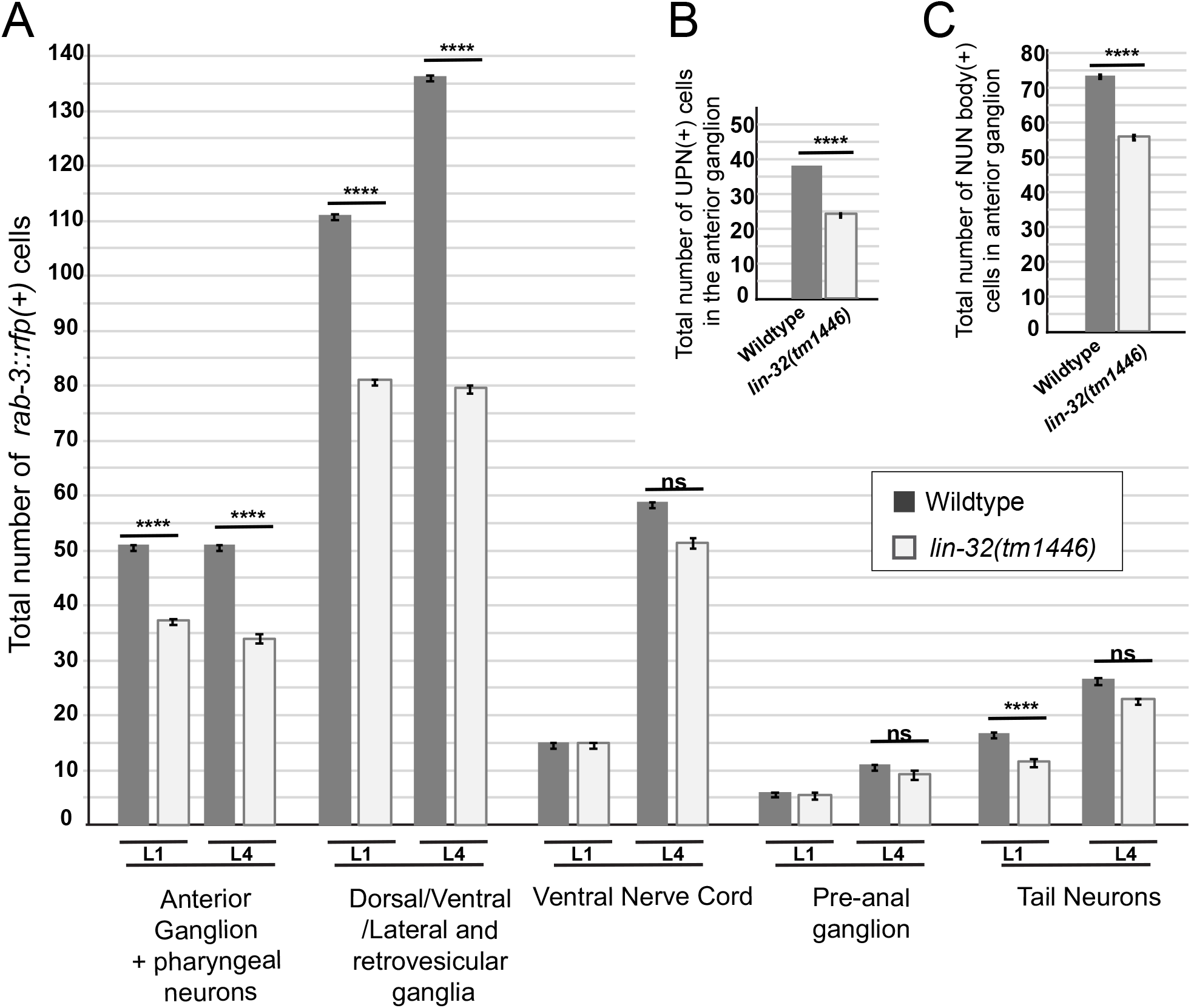
Proneural activities of *lin-32/Ato* throughout the animal. **A:** Expression of the panneuronal gene *rab-3* is affected in different ganglia of the *lin-32/Ato* null allele. The total number of cells expressing the *rab-3* marker *otIs356* are plotted. **B**: The total number of cells expressing the UPN panneuronal marker (contained within the NeuroPAL transgene *otIs669*). The UPN reporter consist of 4 panneuronal promoters from *ric-19, rgef-1, unc-11* and *ehs-1* loci (Yemini *et al*. 2021). We only scored UPN in the anterior ganglion. **C:** The total number of cells with speckled subnuclear morphology (“NUN bodies”) was scored in the anterior ganglion, using Nomarksi optics. The error bars are indicating standard error of mean. Asterisks are indicating significance of p<0.0001. The statistical significance shown were calculated using twoway ANOVA with Tukey test for correction of multiple comparison.

**Fig.3:**
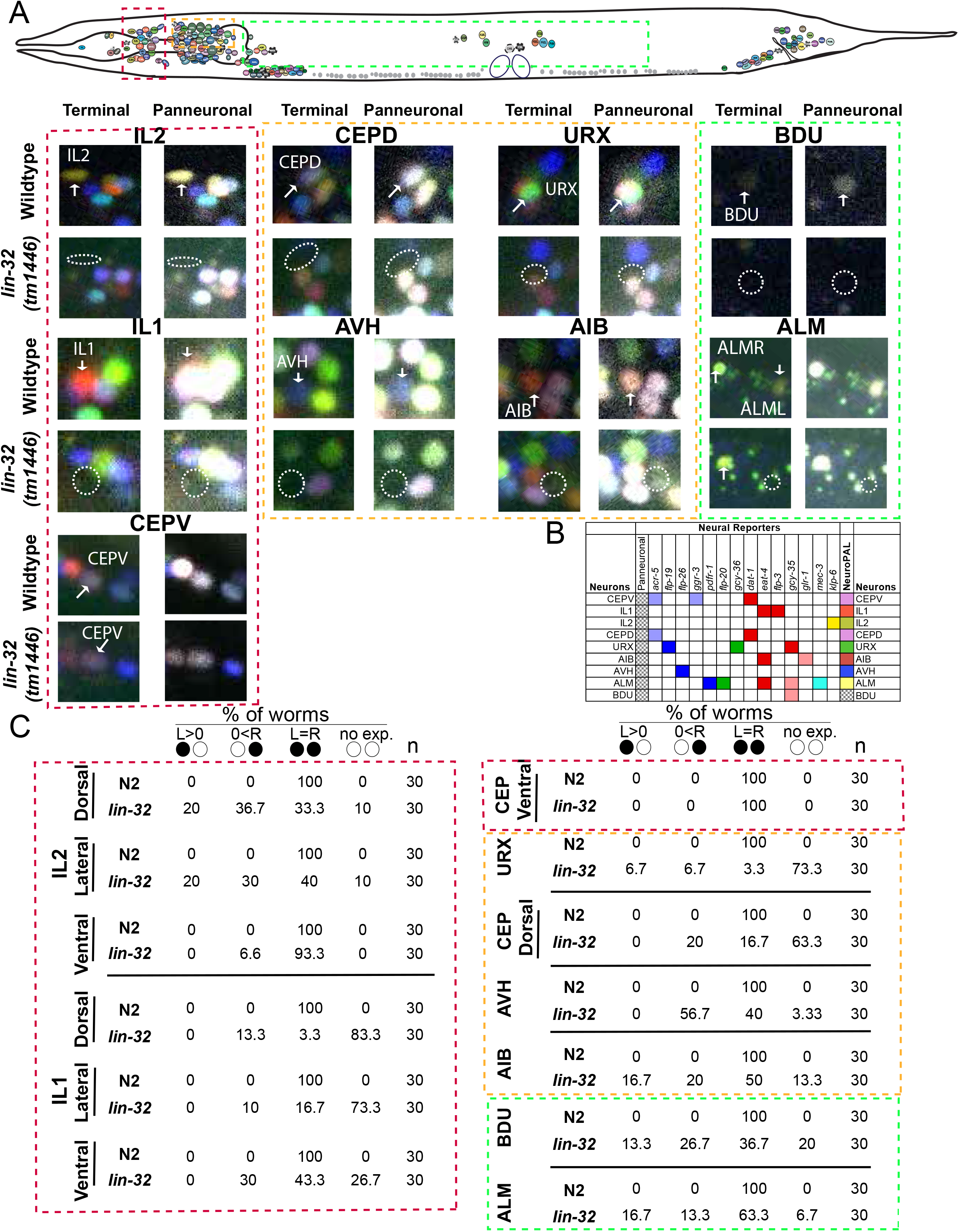
Proneuronal activities of *lin-32/Ato* in specific lineages. **A:** The terminal and panneuronal identities of many neurons that express *lin-32* during development are affected in *lin-32(tm1446)*, as revealed here using the cell-fate marker strain NeuroPAL (*otIs669*). Boxes with different color codes indicate the ganglion for neurons exhibiting fate defects. **B:** The NeuroPAL reporter-fluorophore combinations responsible for distinguishably coloring each neurons type. **C:** Statistics for terminal and panneuronal fate defects shown in panel A. The quantification shown here is the percentage of animals in which the wildtype NeuroPAL color code for the indicated neuron classes is observed in the respective neuron class, as revealed with the NeuroPAL (*otIs669*) transgene. No instances were observed in which the terminal markers were affected while the panneuronal identity remained intact. The same effects of *lin-32/Ato* on *dat-1* expression in CEPD (loss of expression) and CEPV (no effect detected) were also previously reported (Doitsidou *et al*. 2008). Circles indicate bilateral homologs of the respective neuron class (L>0: Expression only in left neuron; 0<R: expression only in right neuron; L=R: expression in both neurons = wildtype).

The hybrid panneuronal marker was expressed from the recently described NeuroPAL transgene, which also contains a large number of additional neuron-type specific markers (Yemini *et al*. 2021). These markers allowed us to more specifically assess which neurons lose their identity and to correlate these losses with normal sites of *lin-32/Ato* expression. We observed panneuronal and cell-type specific marker losses in the AIB and URX neurons, which were previously reported to display differentiation defects in *lin-32* mutants (Rojo Romanos *et al*. 2017; Hori *et al*. 2018)(**Fig.3**). Within the anterior ganglion, we observed losses of panneuronal and cell-specific markers of the radially symmetric IL1 and IL2 sensory neuron classes, consistent with *lin-32/Ato* expression in the lineages that generate these neurons. We independently corroborated the IL1 and IL2 neuron losses with different marker transgenes (*klp-6* for IL2 and *flp-3* for IL1)(**Suppl. Fig.S1**).

In a previous screen for mutants that affect dopaminergic cell fate, we had identified alleles of *lin-32/Ato* and reported that *lin-32/Ato* affects the expression of the dopaminergic marker *dat-1::gfp* in the CEPD neurons (Doitsidou *et al*. 2008). Here, we found that *lin-32/Ato* is expressed in the lineage that gives rise to the CEPD neurons. Using NeuroPAL, we show that *lin-32/Ato* affects expression not only of the *dat-1* marker, but of other cell-identity markers as well, in addition to panneuronal gene expression in CEPD. These results indicate that *lin-32/Ato* acts a proneural factor in this lineage as well (**Fig.3**).

### *lin-32/Ato* is required for terminal selector expression

To further examine the nature of these differentiation defects, we asked whether *lin-32/Ato* controls the expression of the terminal selector-type transcription factors known to specify the identity of the neurons affected by *lin-32/Ato*. We found that the expression of *unc-86*, the identity regulator of the IL2 neurons and the URX neurons is indeed lost in *lin-32/Ato* mutants (**Fig.4**). Expression of *ceh-43/Dlx* and *ceh-32/Six*, candidate terminal selectors for the IL1 neurons, and expression of *lin-11/Lhx1*, a candidate terminal selector for AVJ, are affected in the IL1 and AVJ neurons, respectively (**Fig.4**). Similarly, expression of *ceh-43*, the terminal selector for CEPD neurons (Doitsidou *et al*. 2013) is lost in CEPD of *lin-32/Ato* mutants (**Fig.4**), consistent with the loss of terminal CEPD identity markers in *lin-32* mutants (**Fig.3**)(Doitsidou *et al*. 2008).

**Fig.4:**
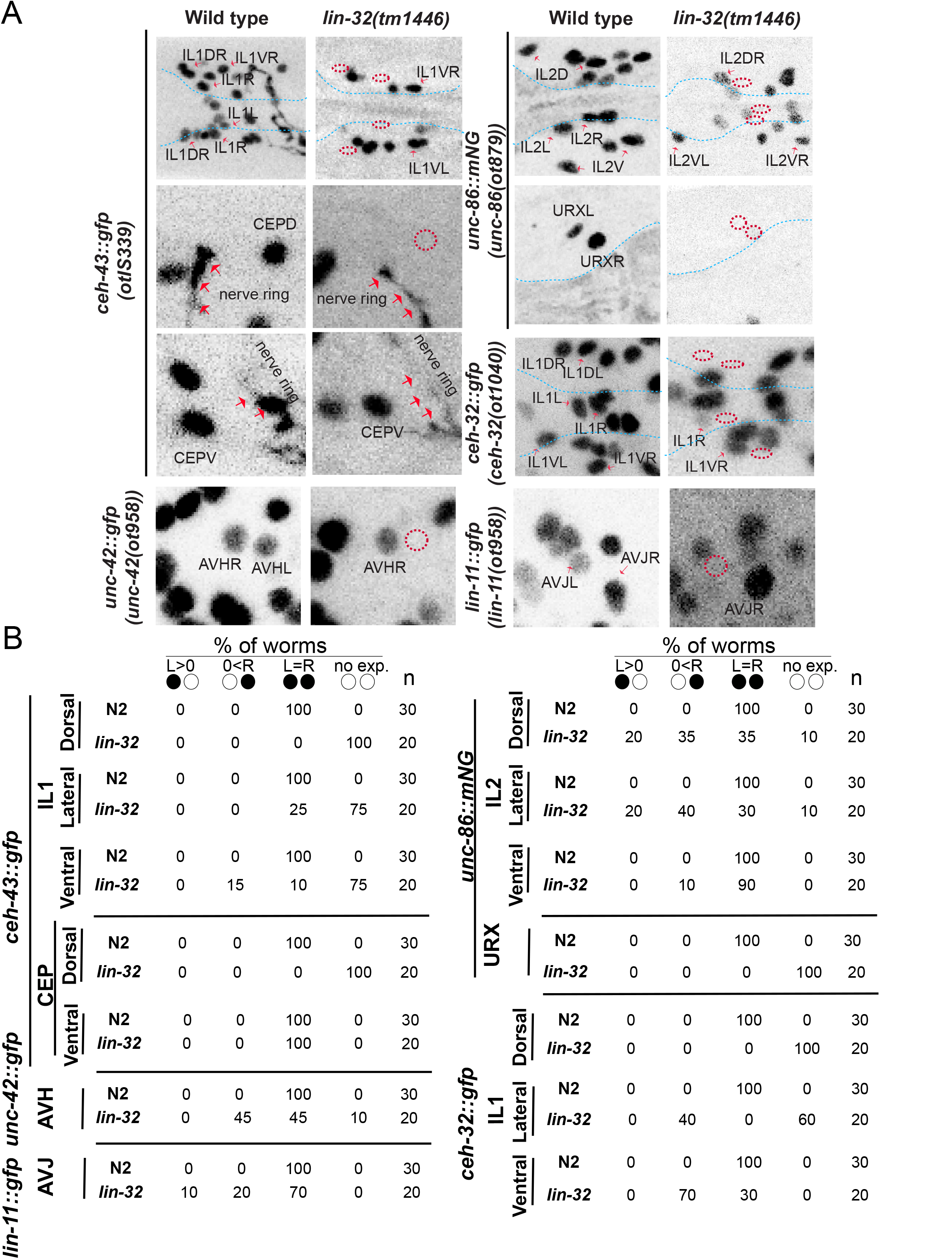
Effect of *lin-32/Ato* on the expression of candidate terminal selectors. **A,B:** Loss of candidate terminal selector expression in *lin-32(tm1446)* null mutants. Reporters are *lin-11(ot958[lin-11::gfp::FLAG], ceh-32(ot1040[ceh-32::gfp]), unc-86(ot879[unc-86::nNeonGreen]), otIs339 (ceh-43^fosmid^::gfp)* and *unc-42 (ot958[unc-42::gfp])* **A:** representative image; **B:** quantification. Circles indicate bilateral homologs of the respective neuron class (L>0: Expression only in left neuron; 0<R: expression only in right neuron; L=R: expression in both neurons = wildtype).

Since terminal selectors like *unc-86, unc-42* and *ceh-43* do not affect panneuronal gene expression (Doitsidou *et al*. 2013; Hobert 2016; Berghoff *et al*. 2021), we conclude that *lin-32/Ato* independently regulates two aspects of the neuronal differentiation programs of cells like the IL2, IL1 or AIB neurons: panneuronal identity and the acquisition of neuron-type specific features via regulation of terminal-selector transcription factors.

### *lin-32/Ato* affects terminal identity markers in subsets of neuronal class members

The effect of *lin-32/Ato* on a number of different neuron classes reveals an interesting phenomenon. All six IL2 class members are very similar neurons based on process projection patterns, synaptic connectivity (White *et al*. 1986) and molecular markers (Taylor *et al*. 2020) and their common identity is specified by the terminal selectors *unc-86, sox-2* and *cfi-1* (Shaham and Bargmann 2002; Zhang *et al*. 2014; Vidal *et al*. 2015). However, we found that the *lin-32/Ato* mutant affects the differentiation of the dorsal and lateral IL2 pairs much more strongly than the ventral IL2 pairs (**Fig.3, Fig.4**). The subclass selective effect of *lin-32/Ato* correlates with the intriguing phenomenon that all six IL2 neurons derive from distinct lineages yet converge on the same neuron type. The dorsal and lateral IL2 pairs are all generated within the ABala lineage branch, while the ventral pairs are generated by the lineally distal ABalpp and ABarap branch (**Fig.1**). In the ABala branch, where loss of *lin-32/Ato* shows an effect, *lin-32/Ato* is normally expressed early in the lineage, while in the branches that produce the ventral pairs in a *lin-32/Ato*-independent manner, *lin-32/Ato* is expressed much later (**Fig.1**).

Another case of subclass expression is observed in the dopaminergic CEP neuron class, composed of a dorsal pair (CEPDL/R) and a ventral pair (CEPDVL/R). These two pairs are anatomically very similar (White *et al*. 1986) and, in the terminally differentiated state, molecularly indistinguishable (Taylor *et al*. 2020). However, each pair derives from distinct embryonic neuroblasts (**Fig.1**). As described above, we found that *lin-32/Ato* has a proneural function in CEPD (**Fig.3)**. However, even though *lin-32/Ato* is expressed in the CEPV neurons (albeit much later than in the CEPD neuron-producing lineage; **Fig.1**), we detected no defect in the generation or differentiation of the CEPV neurons in *lin-32/Ato* null mutants. Panneuronal marker expression and terminal identity markers are unaffected (**Fig.3)** and expression of the terminal selector *ceh-43* is also unaffected (**Fig.4**).

We observed another even more striking case of *lin-32/Ato* selectively affecting on lineally convergent neuron class members, relating to left/right symmetric neuron class members. Within the ABala lineage branch, the left and right AVH neurons are generated from two non-symmetric precursor cells, ABalapaaa (giving rise to AVHL) and ABalappap (giving rise to AVHR)(**Fig.5A**). Another neuron pair generated by these two different blast cells are CANL and CANR (**Fig.5A**). Like the six IL2 neurons, these two left/right symmetric neurons pairs within the ABala lineage are again examples of lineage convergence, where non bilaterally symmetric lineage histories funnel into the generation of indistinguishable left/right neuron pairs. We observed that *lin-32/Ato* is expressed in a left/right asymmetric manner in the lineages that gives rise to AVH and CAN neurons. In the lineage branch that gives rise to the left AVH and the left CAN, *lin-32/Ato* is expressed throughout the lineage, while *lin-32/Ato* is expressed only very late in the postmitotic AVHR and not at all in CANR or the lineage that give rise to it (**Fig.5**). As stated above, we observed that *lin-32/Ato* does not affect the cellular cleavage pattern that gives rise to these cell types. However, we discovered that loss of *lin-32/Ato* affects, in a left/right asymmetric manner, the expression of terminal selector type transcription factors that define the molecular identity of neuron pairs within the ABala lineage. Specifically, *lin-32/Ato* affects expression of *hlh-34*, an identity regulator of the AVH neuron (Berghoff *et al*. 2021), in AVHL, where *lin-32/Ato* expression starts as early as the great-grandmother of AVHL, but not in AVHR, where we observe no *lin-32/Ato* expression (**Fig.5B**).

**Fig.5:**
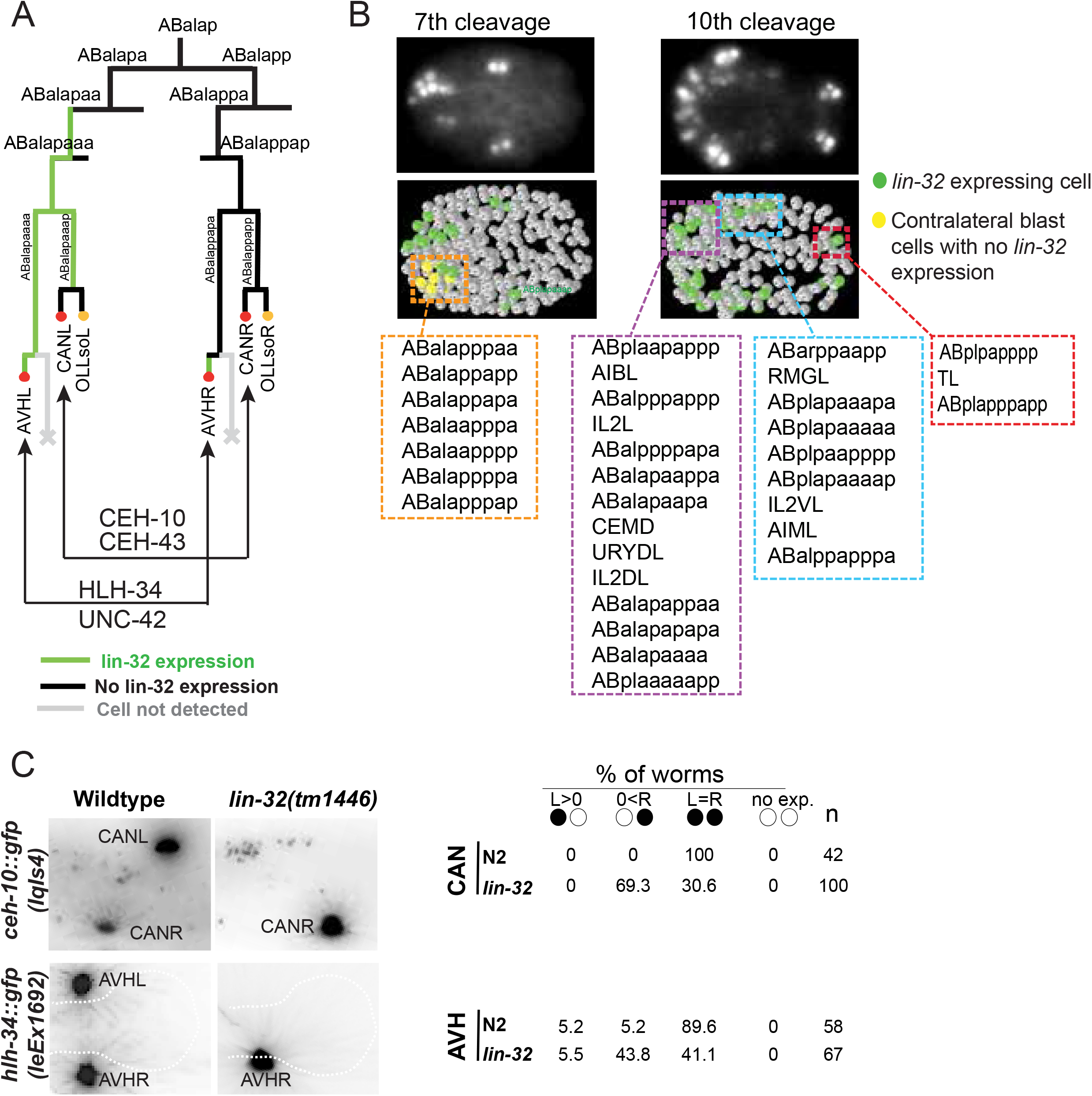
Left/right asymmetric neuronal defects in *lin-32* mutants. **A:** ABalapa descendants are the very first blast cells to show expression of *lin-32*. The ABalapa lineage gives birth to two neurons, AVHL and CANL, among many other neurons. Conversely, their right contralateral homologs derive from the ABalapp lineage, which does not express *lin-32/Ato*. **B:** Representative images of *lin-32/Ato* gene expression at embryonic stages displayed next to their exact time point during embryonic development: the 7^th^ (mitotically active cells) and 10^th^ (postmitotic cells) cleavage. The lower panel shows the ball model for the same embryonic stages. Color-coded boxes represent different regions of the embryo. The name of terminal and blast cells are indicated in the color-coded boxes. **C:** The effect of *lin-32(tm1446)* was quantified using terminal selector reporters for AVH and CAN – *lin-32/Ato* is asymmetrically expressed in their ancestors.

The same left/right symmetric effect is observed on the CAN neuron pair. CAN identity is specified by the Prd-type homeobox gene (Forrester *et al*. 1998; Wenick and Hobert 2004) and may operate together with the Distalless ortholog *ceh-43* (Reilly et al. 2020). We find that *lin-32/Ato* affects expression of the CAN identity regulator *ceh-10* as well as its candidate cofactor *ceh-43/Dlx* in CANL, but not CANR (**Fig.5B**).

### The achaete-scute homolog *hlh-14* provides a mirror image of *lin-32/Ato* function

We considered the possibility that another bHLH transcription factor may provide a mirror-image function of *lin-32/Ato* in controlling the identity of neuron that are contralateral to those affected by *lin-32/Ato*. Unlike *Drosophila* or vertebrates, *C. elegans* only codes for a single Atonal ortholog (Zhao and Emmons 1995; Baker and Brown 2018). The next closely related bHLH genes are *ngn-1* and *cnd-1*, the single Neurogenin and NeuroD orthologs of *C. elegans* which, together with Atonal, form the Ato superfamily of bHLH genes (Hallam *et al*. 2000; Hassan and Bellen 2000; Nakano *et al*. 2010). We generated a strain with a fosmid reporter transgene for *cnd-1*. We also used a previously described *ngn-1* reporter transgene that contains the entire *ngn-1* locus and is capable of rescuing *ngn-1* mutant phenotypes (Nakano *et al*. 2010). These two strains were used to examine *cnd-1* and *ngn-1* expression throughout all stages of embryonic and postembryonic development of the hermaphrodite. We again focused on the AB and MS lineages which produce all but 2 of the 118 neuron classes. We observed expression of both genes in several sublineages, with *ngn-1* being more widespread than *cnd-1* (**Suppl. Fig.S2**). *ngn-1* and *cnd-1* expression is largely nonoverlapping, with the exception of the ABarapp lineage where the expression of both genes overlaps (**Suppl. Fig.S2**). However, we observed no asymmetric expression within bilaterally symmetric neuron pairs in the ABala lineage that would mirror asymmetric expression of *lin-32*.

As the next gene candidates we considered the expression patterns of four homologs of the *Drosophila* Achaete-Scute complex (AS-C): *hlh-6, hlh-19, hlh-3* and *hlh-14*. Their expression patterns had not previously been reported throughout the embryonic nervous system. A fosmid-based *hlh-6* reporter is exclusively expressed in pharyngeal gland cells, as previously reported with smaller reporters (Smit *et al*. 2008). We used CRISPR/Cas9 to tag endogenous *hlh-19* with *gfp* and observed no expression at any stage of embryonic development. In contrast, an *hlh-3* fosmid-based reporter, as well as a CRISPR-generated reporter allele (kindly provided by N. Flames), showed widespread expression throughout the developing embryonic nervous system (**Fig.6**). A *hlh-14* fosmid reporter also showed embryonic expression in neuronal lineages, but its expression was more restricted (**Fig.6**). Intriguingly, in the context of the ABalap lineage where we observed left/right asymmetric *lin-32/Ato* expression (**Fig.1**), we observed that *hlh-14* (but not *hlh-3*) displayed a mirror image asymmetry of this expression. In those left/right symmetric lineages where we observed differential later expression of *lin-32/Ato*, we observed differential earlier expression of *hlh-14*, and vice versa (**Fig.7A-C**).

**Fig.6:**
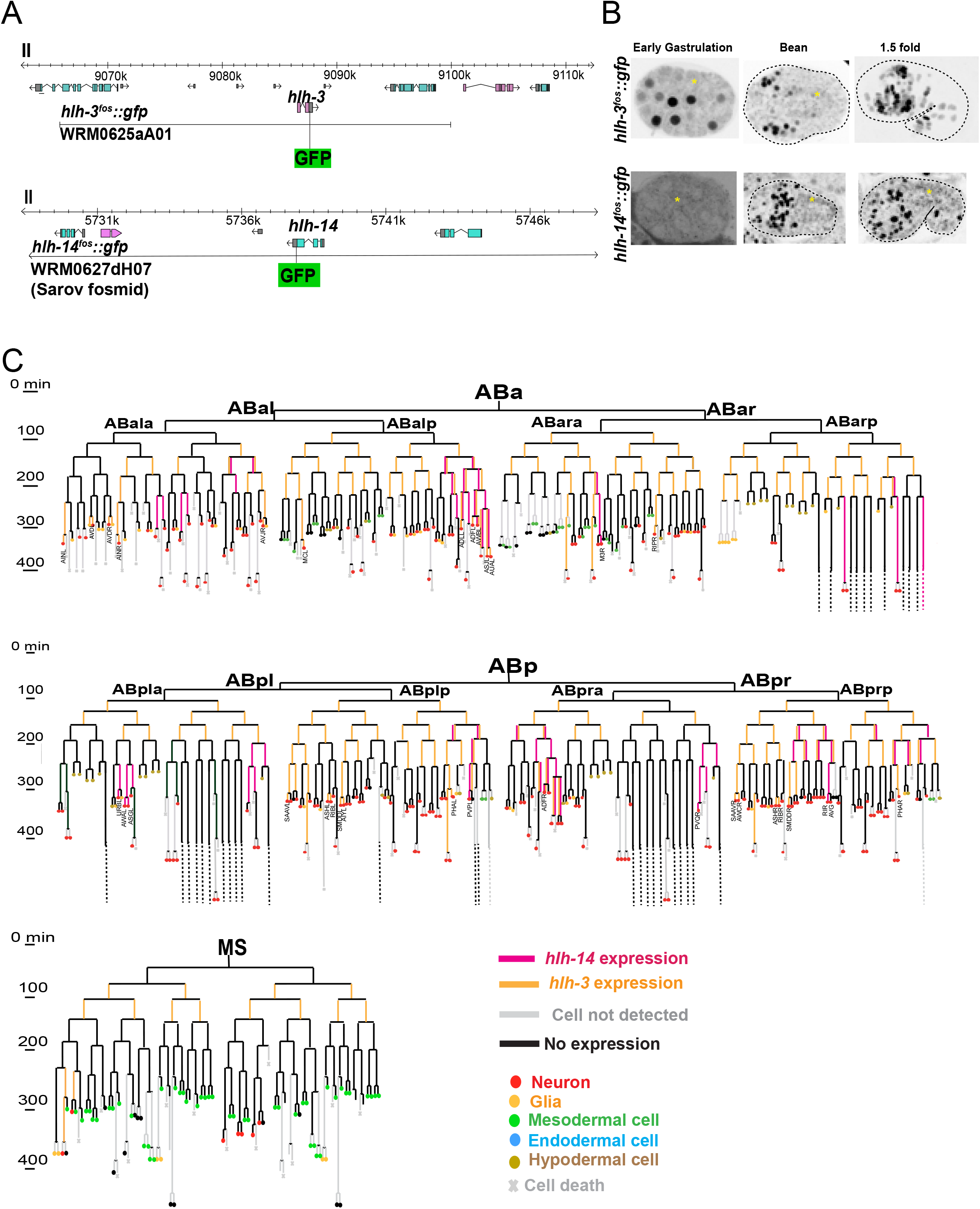
Embryonic expression of the AS-C homologs *hlh-3* and *hlh-14 hlh-3* and *hlh-14* fosmid reporters (*otIs648* and *otIs713*, respectively) and their expression patterns. **A:** Schematic of gene structure and fosmid reporters used for expression pattern analysis. **B:** Representative images shown here are from *hlh-3 and hlh-14* fosmid reporters during different stages of embryonic development; showing the time when the expression starts in blast cells to the time when all postmitotic neurons are born. Yellow asterisks are marking the cytoplasmic autofluorescence. **C:** Lineage diagram showing the expression pattern of *hlh-3* and *hlh-14* during embryogenesis. Our analysis corroborates and extends the previously published expression patterns of *hlh-14* (Frank *et al*. 2003; Poole *et al*. 2011).

**Fig.7:**
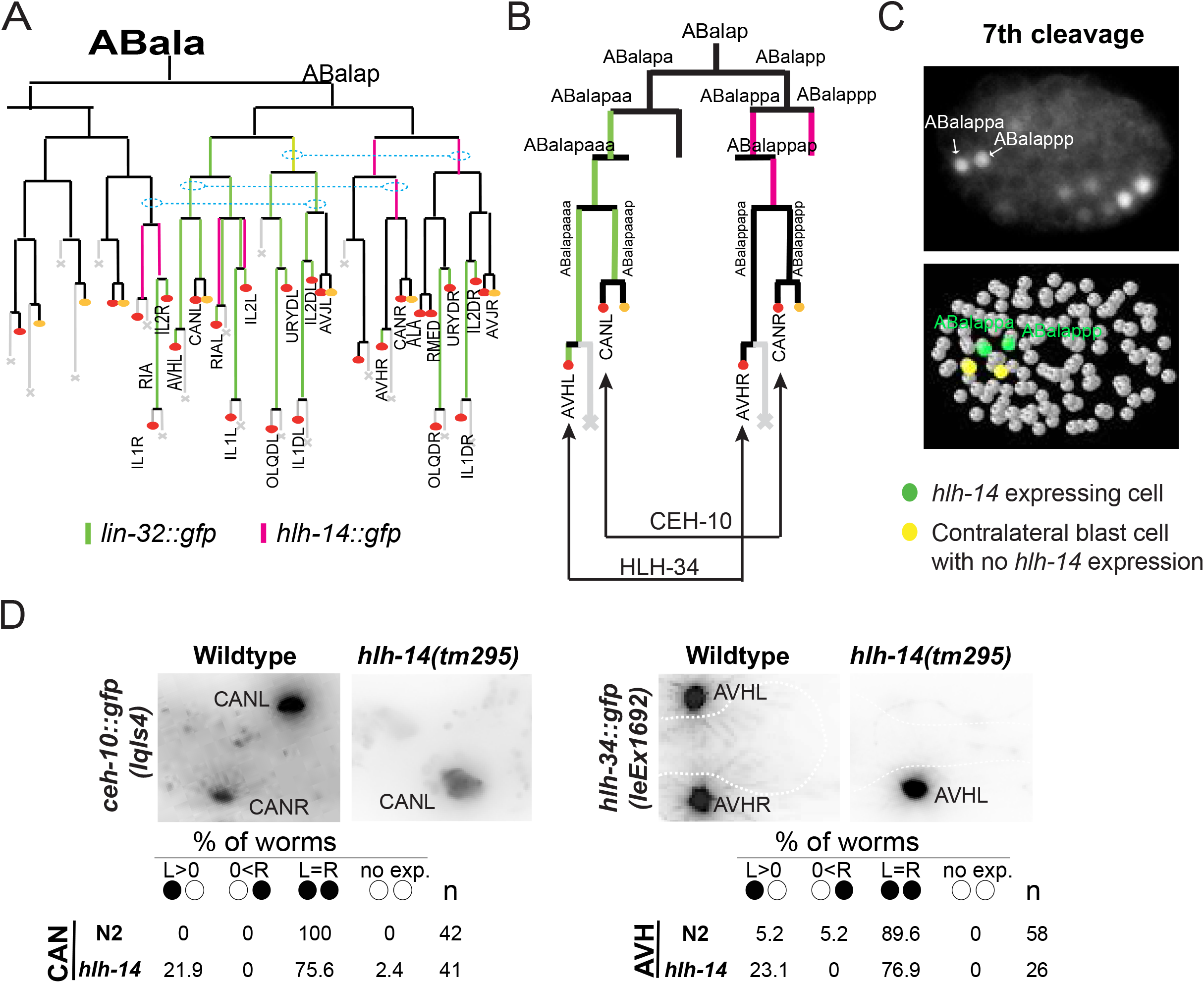
Left/right asymmetric neuronal defects in *hlh-14* mutants. **A:** The lineage diagram shows the expression of *lin-32* and *hlh-14* fosmid reporters in the ABala lineage side by side. *hlh-14* mirrors the expression of *lin-32* in ABalapa descendants, in ABalapp lineage. The dashed lines indicate sublineages that divide in a symmetric manner to produce symmetric cell fates, as shown in panel B. **B:** A more focused version of the diagram in panel A, showing that *hlh-14* expresses in the ABalapp descendants which give birth to AVHR and CANR, a mirror image of *lin-32* expression (Fig.5). **C:** Representative image of *hlh-14* gene expression at the 7^th^ cleavage where mitotically active blast cells ABalappa/p (the great grandmother of AVHR and CANR) start expressing *hlh-14*. The right panel shows the ball model for the same embryonic stages. **D:** The effect of *hlh-14(tm295)* were quantified for terminal selector reporters of AVH and CAN – as was done for *lin-32* mutants (Fig.5C). The wildtype images are from Fig 5C and are shown here for comparison only.

To assess the functional significance of this expression we examined two different left/right symmetric neuron pairs, CANL/R and AVHL/R in *hlh-14* null mutants. We observed mirror image defects in *hlh-14* mutants: CANR, but not CANL, loses *ceh-10* expression (the opposite phenotype as *lin-32/Ato*) and AVHR, but not AVHL, loses *hlh-34* expression (also the opposite phenotype as *lin-32/Ato)(**Fig.7D***). We further note that *hlh-14* is also expressed as a mirror image in other lineages, particularly the IL1 and IL2 lineages, where *lin-32/Ato* shows differential expression in individual class members.

### Neuronal identity transformations in *lin-32/Ato* mutants

Returning to our original analysis of *lin-32/Ato* mutants, we considered cases where we observed expression of *lin-32/Ato*, but no apparent loss of neuronal identity upon loss of *lin-32*. One example is the anterior deirid lineage which produces a group of five bilaterally symmetric neuron pairs (ADE, ADA, AIZ, FLP, RMG) and expresses *lin-32/Ato* early and uniformly (**Fig.8A**). We detected no proneural functions of *lin-32/Ato* in this lineage (*i.e*. no loss of the panneuronal marker) and we also observed no obvious defects of the cellular cleavage pattern in the lineage. A marker (*dat-1::gfp*) expressed in of one neuron class in the lineage, the dopaminergic ADE neuron, is expressed in several additional cells in *lin-32/Ato* mutants (Doitsidou *et al*. 2008), but it had been unclear whether these cells are ectopically generated cells or whether the marker is aberrantly expressed in other cells of this lineage. The use of NeuroPAL (Yemini *et al*. 2021), which provides unique labels to all cells in the lineage, provided us with the opportunity to assess this defect in more detail. We independently confirmed the ectopic dopamine marker gene expression and found that it is paralleled by a loss of expression of markers for the AIZ, ADA, FLP and RMG neurons (**Fig.8B,D**). Hence, AIZ, ADA, FLP and RMG appear to have transformed their identity to that of the ADE neurons.

**Fig.8:**
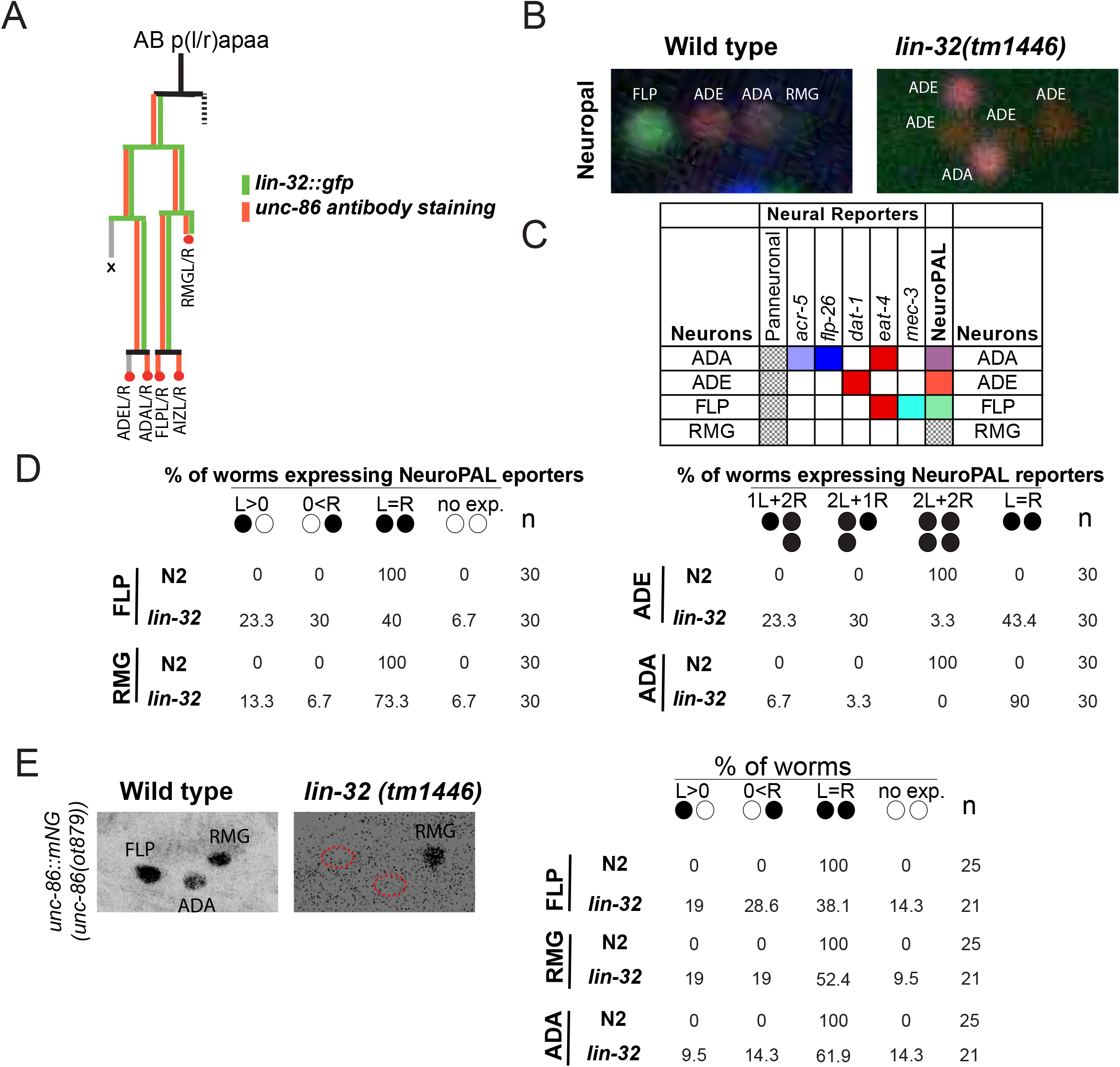
*lin-32/Ato* mutants display neuronal identity fate transformation in anterior deirid lineage. **A:** The expression of *lin-32/Ato* is shown in lineages giving birth to the anterior deirid neurons. The expression of *lin-32/Ato* overlaps perfectly with the previously reported *unc-86* expression (Finney and Ruvkun 1990). **B:** The neuronal identities of Anterior deirid neurons were observed using NeuroPAL *otIs669*. In *lin-32(tm1446)* mutants there are ectopic neurons expressing ADE-specific terminal markers while the terminal markers for FLP, RMG and sometimes the ADA neurons are lost. Quantification is shown in panel D. **C:** The table shows the combination of different reporters in NeuroPAL strain used to mark cells in this lineage. **D:** This panel shows the quantification of different phenotypic categories observed in *lin-32(tm1446)* as compared to wildtype using NeuroPAL reporters. Circles indicate bilateral homologs of the respective neuron class (L>0: Expression only in left neuron; 0<R: expression only in right neuron; L=R: expression in both neurons = wildtype; 2R or 2L: expression in an additional neuron on left or right). **E:** *unc-86* expression (reporter allele *ot879*) is lost in the FLP, ADE and RMG neurons in the *lin-32* mutants. Quantification is shown in right panel.

All of the 4 neuron classes that transform to ADE identity in *lin-32/Ato* mutants, normally express the *unc-86/Brn3* POU homeobox gene (Finney and Ruvkun 1990; Serrano-Saiz *et al*. 2018)(**Fig.8A**). *unc-86/Brn3* acts as terminal selector in at least one of these neurons, FLP (Topalidou and Chalfie 2011), and perhaps others as well (Serrano-Saiz *et al*. 2013). We therefore tested whether loss of *lin-32/Ato* function affects *unc-86* expression in these cells. Using a *mNeonGreen-tagged unc-86* locus as a reporter for *unc-86* expression (Serrano-Saiz *et al*. 2018), we indeed observed a loss of *unc-86* expression in all 4 neuron classes of the lineage (**Fig.8E**). A similar result was also described previously using an antimorphic allele of *lin-32/Ato, u282* (Baumeister *et al*. 1996). Future work will determine whether the neuronal identity transformation observed in *lin-32/Ato* mutants in the anterior lineage is entirely explicable through the loss of *unc-86* expression.

We observed another cell identity transformation in *lin-32/Ato* mutant animals in lineages that generate the four radially symmetric OLQ and URY neurons (Fig.9A). These neurons derive from four neuroblasts: ABalapapap, ABalapppap, and the two bilaterally symmetric ABp(l/r)paaappp neurons. All of these neuroblasts were *lin-32/Ato* positive and maintained *lin-32/Ato* expression throughout ensuing divisions. As in the anterior deirid lineage, we found no proneural functions of *lin-32/Ato* in these lineages, *i.e*. we observed no obvious defects in cellular cleavage pattern in the lineage until the 300-minute stage (Supp Dataset 1), nor did we observe loss of panneuronal reporter expression. Instead, we observed another apparent neuronal identity transformation: OLQ neurons lose characteristic marker gene expression and instead gain expression of URY marker genes (Fig.9B-E). As in the anterior lineage, we sought to extend this observation by analyzing the expression of potential cell-identity regulators in this lineage. A potential URY identity regulator is the homeobox gene *ceh-32/Six3*, which is expressed in URY but not OLQ (Reilly *et al*. 2020). We observed that in *lin-32/Ato* mutants, OLQ neurons gain *ceh-32* expression (Fig.9F). Taken together, we conclude that *lin-32/Ato*, either directly or indirectly, regulates the expression of terminal selector-type transcription factors that play a role in promoting specific neuronal identities and suppressing alternative identities.

**Fig.9:**
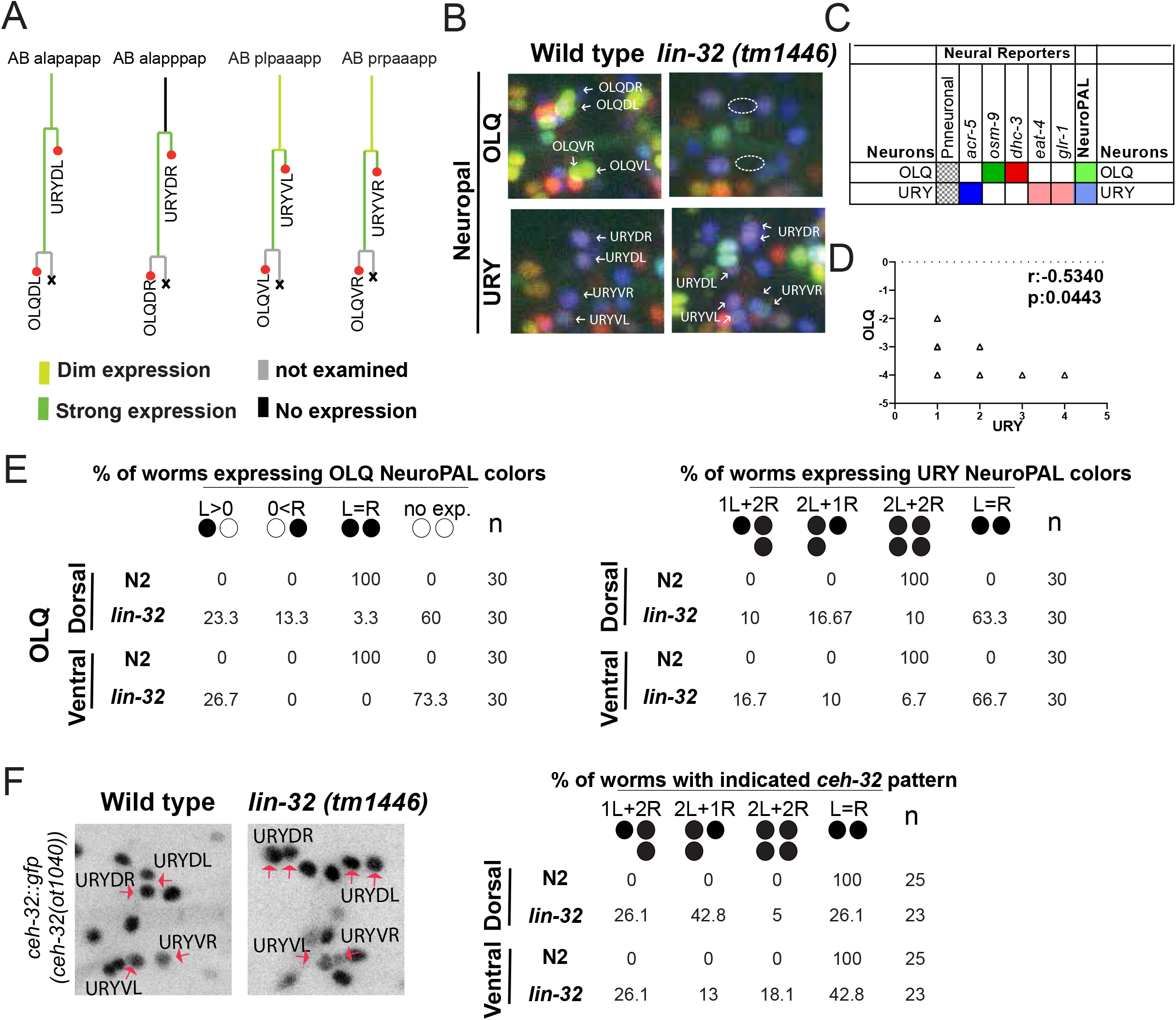
OLQ to URY identity transformation in *lin-32/Ato* null mutants. **A:** The lineage diagrams show the expression of *lin-32/Ato* in lineages giving birth to URY and OLQ neurons. **B:** The terminal identity of OLQ and URY neurons is marked by NeuroPAL (*otIs669*). In *lin-32(tm1446)* mutants there are ectopic signals for the terminal markers of URY while the OLQ specific signals were lost. Specifically, we observed potential duplicates of URYDR and URYVR in these mutants. Quantification is shown in panel I. **C:** The table shows the combination of different reporters mixed in NeuroPAL strain used to mark URY and OLQ neurons. **D:** Correlation between the gain in URY-terminal markers and the loss of OLQ-terminal markers. URY and OLQ are lineal cousins of each other and their gain-loss correlation is statistically significant as tested by Pearson correlation (r=-0.534). The negative correlation value reflects the anticorrelated relationship between the observation of both types of neuron, meaning that observing URY is correlated with not observing its lineal cousin OLQ. **E:** Quantification of NeuroPAL color codes observed in *lin-32(tm1446)* compared to the wildtype, using the NeuroPAL reporter (*otIs669*). Left panels shows OLQ color code in ventral and dorsal OLQ neurons; this color code is partially lost in *lin-32* mutants. Right panels shows URY color codes, which are observed in additional cells in *lin-32* mutants. Circles indicate bilateral homologs of the respective neuron class (L>0: Expression only in left neuron; 0<R: expression only in right neuron; L=R: expression in both neurons = wildtype; 2R or 2L: expression in an additional neuron on left or right). **F:** Ectopic OLQ expression of the *ceh-32* reporter allele (*ot1040*), normally expressed in URY. Quantification is shown in right panel.

## DISCUSSION

We found that, as expected from its previously described function in postembryonic development (Zhao and Emmons 1995), *lin-32/Ato* acts as a proneural gene in a number of embryonically generated lineages. One distinguishing feature of the proneural role of *lin-32/Ato* in postembryonic versus embryonic development is that, in postembryonic development, a loss of the proneural activity of *lin-32/Ato* results in a termination of neuroblast divisions and conversion of the neuroblast to that of an ectodermal skin cells (Zhao and Emmons 1995). In contrast, during embryogenesis, we found no evidence that the loss of *lin-32/Ato* affects cellular cleavage patterns or results in obvious conversions to hypodermal fate. What we do observe is a loss of expression of panneuronal features, a loss of cell identity-controlling transcription factors and, consequently, lost expression of cell-specific identity features.

One intriguing aspect of the proneural function of *lin-32/Ato* relates to the highly selective effect that *lin-32/Ato* has on specific members of individual neuron classes. This observation provides a molecular correlate to the phenomenon of lineage convergence: cells with different lineage histories acquire similar terminal identities. Lineage convergence is widespread in the *C. elegans* nervous system (Sulston *et al*. 1983) and, with the advent of novel means to lineage more complex organisms, has now been observed in vertebrates as well (Mckenna *et al*. 2016; Wagner *et al*. 2018; Cao *et al*. 2019; Chan *et al*. 2019). In the *C. elegans* nervous system, divergent lineages of individual neuron class members converge on similar gene expression profiles via the activation of terminal selectors of neuronal identity in individual class members. For example, the six IL2 sensory neurons arise from different, non-homologous lineages but are all specified by the combinatorial activity of three terminal selectors, *unc-86, cfi-1* and *sox-2* (Shaham and Bargmann 2002; Zhang *et al*. 2014; Vidal *et al*. 2015). Likewise, the lineally diverse CEP neuron class members are all specified by the combinatorial activity of the terminal selectors *ast-1, ceh-43* and *ceh-20* (Flames and Hobert 2009; Doitsidou *et al*. 2013). We have shown here that different class members are specified by distinct upstream inputs. Specifically, *lin-32/Ato* controls the generation of distinct subsets of members from a given neuron class. In the most extreme cases, we observed that the left neuron of indistinguishable bilaterally symmetric neurons classes is controlled by *lin-32/Ato*. Intriguingly, the right neuron of these neuron classes is controlled by a distinct bHLH gene, *hlh-14*. Although we have not taken our study here to the *cis*-regulatory level, it is conceivable that the *cis*-regulatory control regions of terminal selectors serve as “integration devices” that sample distinct lineage inputs.

Apart from proneural functions in a number of different cellular context, we also identified roles of *lin-32/Ato* in distinguishing the execution of distinct neuronal differentiation programs, such that the loss of *lin-32/Ato* leads to a conversion of one neuronal fate in that of another neuron fate. We observed such homeotic identity transformations in multiple distinct lineal contexts and we propose that these are also the result of lost terminal selector expression. Previous work has shown that in multiple different cellular contexts, terminal selectors can act in a mutually antagonistic and competitive manner, such that removal of one selector may now enable a different selector to exert its function (Arlotta and Hobert 2015). For example, loss of the *mec-3* homeobox gene in the ALM neuron allows UNC-86 to pair up with a different transcription factor, PAG-3, to now promote BDU neuron fate in the ALM neuron (Way and Chalfie 1988; Gordon and Hobert 2015). In analogy to these cases, it is conceivable that the lost expression of some terminal selectors in *lin-32/Ato* allows other terminal selectors to promote alternative fates.

In conclusion, we have provided insights into how terminal differentiation programs in the nervous system, controlled by terminal selector transcription factors, are coupled to earlier developmental events, and, specifically, to the lineage history of a cell. Our findings demonstrate that terminal selectors are key integrators of lineage history and provide novel perspectives on how proneuronal genes function to pattern the nervous system.

## ACKNOWLEDGMENTS

We thank Chi Chen for assistance with microinjections to generate strains, Cyril Cros, Maryam Majeed, Emily Berghoff, Nuria Flames, Erik Lundquist, Sarah Finkelstein, and Hana Littleford for kindly providing strains, and Iva Greenwald, Richard Poole and members of the Hobert lab for comments on the manuscript. This work was funded by the Howard Hughes Medical Institute.

## SUPPLEMENTARY LEGENDS

**Suppl. Dataset 1**: **Lineage movie of *lin-32/Ato* animals**. Two embryos from *lin-32(tm1446)* were lineaged using the classic Nomarski-based technique (Schnabel *et al*. 1997). The movies are in Laura Waivier format and can be seen using SIMIBiocell software.

**Suppl. Fig.S1:**
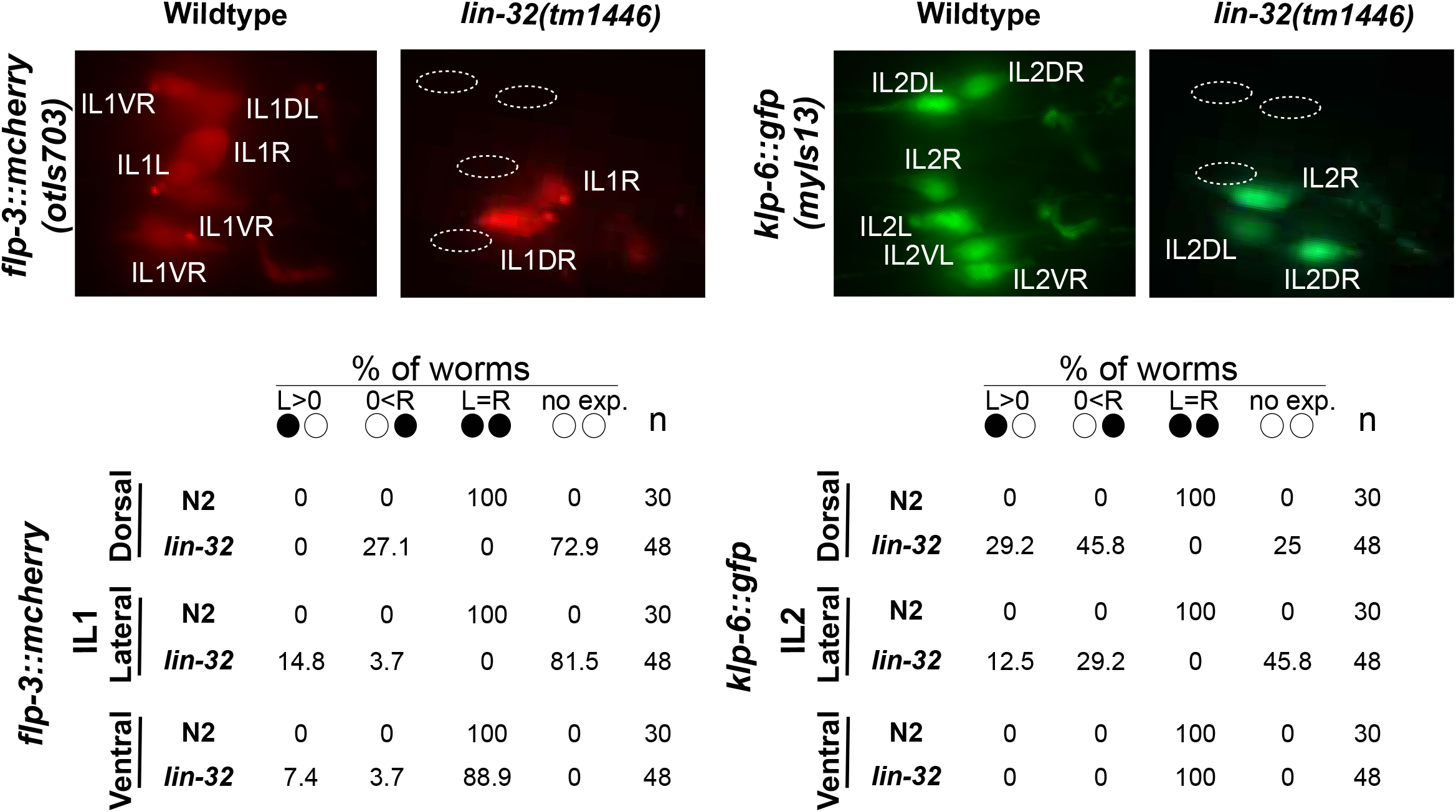
Additional effects of *lin-32/Ato* on marker gene expression. The terminal identity of IL1 and IL2 neurons are affected in *lin-32(tm1446)*, shown here using *flp-3::mCherry* and *klp-6::gfp* reporters. Quantification is shown in lower panels. Circles indicate bilateral homologs of the respective neuron class (L>0: Expression only in left neuron; 0<R: expression only in right neuron; L=R: expression in both neurons = wildtype).

**Suppl. Fig.S2:**
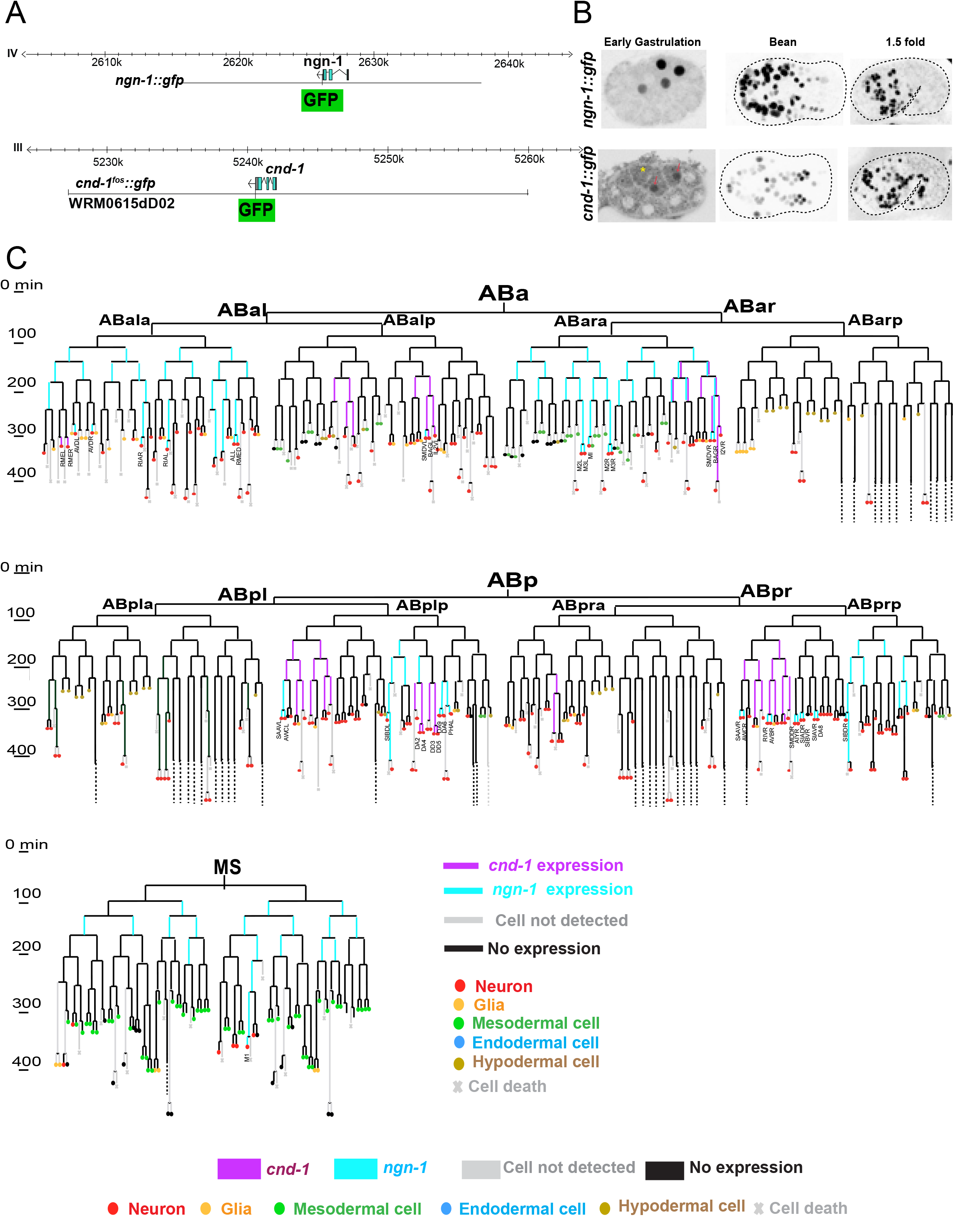
Embryonic expression of the *C. elegans* neurogenin and NeuroD homologs, *ngn-1* and *cnd-1*. *cnd-1 and ngn-1* reporters (*otIs813* and *nIs394*, respectively) and their expression patterns. **A:** Schematic of gene structure and the fosmid used for expression pattern analysis. **B:** Representative images of *cnd-1 and ngn-1* gene expression at embryonic stages, starting from the earliest stage where the expression starts and continuing to the time when all terminal neurons have been born. Yellow asterisk is marking the cytoplasmic autofluorescence and red arrows are pointing toward real expression inside nuclei. **C:** The full lineage of *cnd-1 and ngn-1* expression in the AB and MS lineages; these lineages produce all but 2 of the 302 neurons in adult *C. elegans* hermaphrodites. Our analysis confirms previously published expression patterns reported for a subset of the cells shown here (Hallam *et al*. 2000; Nakano *et al*. 2010).

